# Complex ecological phenotypes on phylogenetic trees: a Markov process model for comparative analysis of multivariate count data

**DOI:** 10.1101/640334

**Authors:** Michael C. Grundler, Daniel L. Rabosky

## Abstract

The evolutionary dynamics of complex ecological traits – including multistate representations of diet, habitat, and behavior – remain poorly understood. Reconstructing the tempo, mode, and historical sequence of transitions involving such traits poses many challenges for comparative biologists, owing to their multidimensional nature. Continuous-time Markov chains (CTMC) are commonly used to model ecological niche evolution on phylogenetic trees but are limited by the assumption that taxa are monomorphic and that states are univariate categorical variables. A necessary first step in the analysis of many complex traits is therefore to categorize species into a pre-determined number of univariate ecological states, but this procedure can lead to distortion and loss of information. This approach also confounds interpretation of state assignments with effects of sampling variation because it does not directly incorporate empirical observations for individual species into the statistical inference model. In this study, we develop a Dirichlet-multinomial framework to model resource use evolution on phylogenetic trees. Our approach is expressly designed to model ecological traits that are multidimensional and to account for uncertainty in state assignments of terminal taxa arising from effects of sampling variation. The method uses multivariate count data for individual species to simultaneously infer the number of ecological states, the proportional utilization of different resources by different states, and the phylogenetic distribution of ecological states among living species and their ancestors. The method is general and may be applied to any data expressible as a set of observational counts from different categories.

---

Most species in the natural world make use of multiple, categorically-distinct types of ecological resources. Many butterfly species use multiple host plants, for example (Ehrlich & Raven 1964 10/7/19 7:40:00 PM; Robinson 1999). Insectivorous warblers in temperate North America use multiple distinct microhabitats and foraging behaviors (MacArthur 1958), as do honeyeaters in mesic and arid Australia (Miller et al. 2017). The evolution of novel patterns of resource use can impact phenotypic evolution (Martin & Wainwright 2011; Davis et al. 2016), diversification (Mitter et al. 1988; Givnish et al. 2014), community assembly (Losos et al. 2003; Gillespie 2004), and ecosystem function (Harmon et al. 2009; Bassar et al. 2010). Consequently, there has been substantial interest in understanding how ecological traits related to resource use evolve and for exploring their impacts on other evolutionary and ecological phenomena (Vrba 1987; Futuyma & Moreno 1988; Forister et al. 2012; Price et al. 2012; Burin et al. 2016).

Making inferences about the evolutionary dynamics of resource use, however, first requires summarizing the complex patterns of variation observed among taxa into traits that can be modeled on phylogenetic trees. Typically, this is done by explicit or implicit projection of a multidimensional ecological phenotype into a greatly simplified univariate categorical space (e.g. “carnivore”, “omnivore”, “herbivore”). It is widely recognized that the real-world complexities of resource use are not adequately described by a set of categorical variables (Hardy & Linder 2005; Hardy 2006). Nonetheless, it is also true that major differences in resource use can sometimes be summed up in a small set of ecological states, a point made by Mitter et al. (1988) in their study of phytophagy and insect diversification. For this reason, continuous-time Markov chain (CTMC) models, which require classifying species into a set of character states, have become commonplace in macroevolutionary studies of ecological trait evolution (Kelley & Farrell 1998; Nosil 2002; Price et al. 2012; Hardy & Otto 2014; Cantalapiedra et al. 2014; Burin et al. 2016). CTMC models describe a stochastic process for evolutionary transitions among a set of character states and are used to infer ancestral states and evolutionary rates, and to perform model-based hypothesis tests (O’Meara 2012). Many complex ecological traits, however, will be poorly characterized by a set of univariate categorical states. Consequently, species classified in single state can nonetheless exhibit substantial differences in patterns of resource use, creating challenges for interpreting evolutionary transitions among character states as well as for understanding links between character state evolution and diversification.

An additional limitation of continuous-time Markov chains for modeling resource use evolution emerges from the fact that species are classified into ecological states without regard for the quality and quantity of information available to perform the classification exercise. As an example, species with few ecological observations might be classified as specialists for a particular resource, when their apparent specialization is strictly a function of the small number of ecological observations available for the taxon. More generally, by failing to use a statistical model for making resource state assignments, we neglect a major source of uncertainty in our data: the uneven and incomplete knowledge of resource use across different taxa. This uncertainty, in turn, has substantial implications for how we project patterns of resource use onto a set of resource states. By failing to account for uneven and finite sample sizes characteristic of empirical data on resource use we cannot be certain if state assignments reflect true similarities or differences in resource use or are merely the expected outcome of sampling variation.

Consider the simple example in Figure 1, consisting of four species and three resources. Panels (a) and (b) illustrate the true resource states and their phylogenetic distribution across a set of four species and their ancestors. Here, an ancestral specialist evolved a generalist diet via a single transition (panel b), such that there are two extant species with the ancestral specialist diet (species X and Y) and two with the derived generalist diet (species P and Q). Panel (c) illustrates the observation process: empirical observations on diet are collected for each taxon, which are influenced by sampling variation. In panel (d), these empirical observations are used to classify each species into one of two diet states, based on the sampled relative importance of the three food resource categories (e.g., panel c). These relative importance estimates are based on uneven, and in some cases quite small, sample sizes, consistent with many empirical datasets (Vitt & Vangilder 1983; Shine 1994; Alencar et al. 2013). In panel (e) we imagine repeating the state assignment process on independent datasets while holding the samples sizes fixed to those in panel (c), which reveals that both the initial state assignments and the number of states from (d) are highly sensitive to real-world levels of sampling variation. This has obvious implications for downstream macroevolutionary analyses. There is a serious risk that incorrect classification, and therefore spurious diet state variation, will emerge from sampling variation alone. In the analyses that underlie Figure 1, we find that more than 70 percent of tip state classifications do not match the true pattern of resource use.

**Figure 1.**
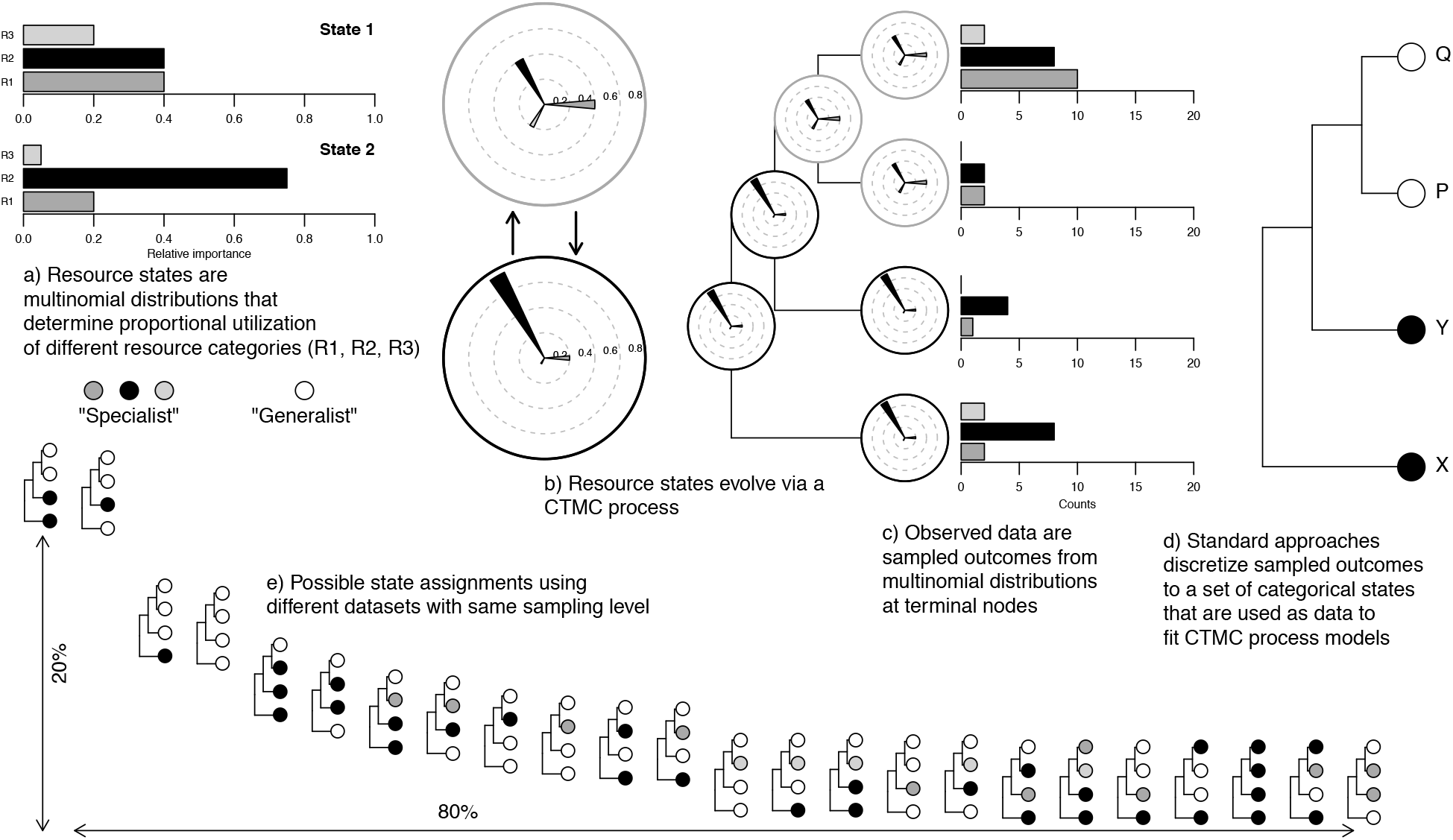
Distribution and representation of multivariate ecological phenotypes (a, b, c), data as sampled by researchers (c), and sampled states as typically represented by univariate categorical traits (d, e). Loss and distortion of information associated with complex phenotypes motivates the development of the Dirichlet-multinomial model described in this article. a) True resource states are multinomial distributions that determine the proportional utilization of three dietary resource categories by four species. Species X and Y are primarily specialists on resource category R2 whereas species P and Q exhibit generalized use of all three resource categories. b) The resource state of a species is the outcome of evolution via a CTMC process where the states correspond to multinomial distributions over a set of resource categories. Here, the multinomial distributions from (a) are represented as rose plots: the direction of a spoke identifies the resource category and the length of a spoke is equal to the proportional utilization of that category. The phylogeny depicts the true evolutionary history of change. c) Sampling: empirical data are sampled outcomes from these multinomial distributions, and the number of observations can differ among species. Here, for example, species Q has 20 observations and species P has just 4. d) Univariate projection: typically, these multivariate outcomes are projected onto univariate categorical resource states, resulting in loss of information and sensitivity to sampling variation. In this example, a species is considered a “specialist” on a particular category if the sampled proportion of the category exceeds 0.7. Otherwise, it is considered a “generalist”. In this case, the dataset and cutoff value align to match each species with its correct modal resource category. e) Simulation illustrating how univariate projection and sampling effects can lead to spurious variation in state assignments and incorrect evolutionary histories. The sampling and projection process illustrated above was repeated 1000 times, holding true resource distributions (a) and sample sizes (c) identical. State assignments are sorted along the x-axis according to their frequency of occurrence in the simulated datasets. Note that the procedure correctly matches all species with their modal food resource in a minority of cases and results in a variable number of states across datasets. The implication for macroevolutionary studies is that we cannot be certain whether state assignments are reflective of true patterns of resource use or are merely the expected outcome of sampling variation and projection.

Is this a problem in practice? This issue is difficult to assess because few studies provide information about the sample sizes that underlie state assignments. In most cases, ecological states are simply asserted as known. It is also important to emphasize that the specific problem in Figure 1 is an outcome of a more general problem: standard CTMC models have a limited ability to model complex ecological phenotypes because of the assumption that states in the model are univariate categorical variables (Hardy & Linder 2005; Hardy 2006). While it is true that CTMC models operate on a countable state space, it is not true that the states of the system must be univariate categorical variables. In this paper we treat states as multivariate probability distributions to develop a model for studying the evolutionary dynamics of ecological resource use on phylogenetic trees that avoids the need to classify taxa into univariate categorical states.

Our approach is explicitly designed to model resource traits that are multidimensional and to account for uncertainty in ecological state assignments of terminal taxa arising from effects of sampling variation. Antecedents to our approach can be found in the use of hidden Markov models in macroevolution (Felsenstein & Churchill 1996; Marazzi et al. 2012; Beaulieu et al. 2013; Beaulieu & O’Meara 2016; Caetano et al. 2018), which assume a deterministic many-to-one mapping from a set of hidden states to a set of observable states. By contrast, we assume that each hidden state is a multinomial distribution and that observed data are sampled outcomes from these distributions (see panels (a) through (c) of Fig. 1). The number of states in the model and the states themselves are not directly observed and are estimated from the data. Using simulations and an empirical dataset of snake diets, we show how the method can use observational counts to simultaneously infer the number of resource states, the proportional utilization of resources by different states, and the phylogenetic distribution of ecological states among living species and their ancestors. The method is general and applicable to any data expressible as a set of observational counts from different resource categories.

## MATERIALS & METHODS

### General overview

We assume there is a discrete set of resource niches, which we refer to as “resource states” or, more simply, as “states”. We use the term resource in a broad sense, with the understanding that it may refer to behavior, habitat, prey, predators, etc. Although the set of resource states is finite, we imagine that a state assigns a taxon a continuously-valued parameter that determines its proportional utilization of different resources. Taxa that share a resource state therefore exhibit similar patterns of resource utilization. The data for each taxon consist of a set of counts, with each count recording the number of times a specific taxon was observed to use a specific resource. We refer to different resources as “resource categories”. The set of resource states and their associated distributions over resource categories are therefore not directly observed but must be estimated from empirical count data. We assume that resource states evolve along a phylogenetic tree according to a CTMC process and that empirical counts are observations drawn from a multinomial distribution associated with each resource state. Our approach is similar to phylogenetic threshold models that combine a probability model for the evolution of an unobserved variable and a probability model for sampling the observed data conditioned on the set of unobserved variables (Felsenstein 2012; Revell 2014). The general approach of treating states as probability distributions is due to Baum and Petrie (1966).

### Model description

Let ***D*** = {***d***_1_,…, ***d**_n_*} denote the data observed for a set of *n* related taxa. Each datum ***d**_i_* = {*d*_*i*1_,…, *d*_*ij*_} records the number of times the *i*-th taxon was observed to use each of ***J*** possible resource categories. For example, in MacArthur’s (1958) study on foraging behaviors in wood-warblers, he made discrete observations of foraging orientation (radial, tangential, vertical) for five species. For Blackburnian warblers, observations (counts) for each of these behaviors were 11, 1, and 3; the resource vector for this species is thus d = {11, 1, 3}, for a total of 15 observations across all three “resource categories”. We assume that each taxon belongs to one of *K* possible states, and we let ***X*** = {*X_1_*,…,*X_n_*}, where each *X_i_* ∈ {1,…,*K*} denotes the state of a particular taxon. States are unobserved but are assumed to represent distinct resource niches so that ***X*** partitions ***D*** into subsets of taxa with homogeneous patterns of resource utilization. Our target of inference is the posterior distribution of resource state assignments *p*(***X***|***D***) ∝ *p*(***D***|***X***)*p*(***X***). To that end we define a probability model for the data as

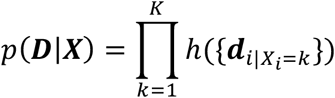

where *h*({***d***_*i*|*X_i_*=*k*_}) is the marginal likelihood function of the data for taxa assigned to state *k*. We assume that

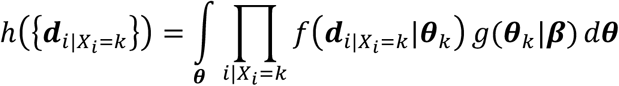

where *f* is a multinomial density with parameter ***θ***_*k*_ and *g* is a Dirichlet prior on ***θ***_*k*_ with hyperparameter ***β***. Because the Dirichlet distribution is conjugate to the multinomial distribution this parameterization allows us to analytically marginalize over the unknown multinomial parameters. Concretely, this means that *h*({***d***_*i*|*X_i_*=*k*_}) is a Dirichlet-multinomial density, which, ignoring multinomial coefficients associated with each ***d***_*i*_, is equal to

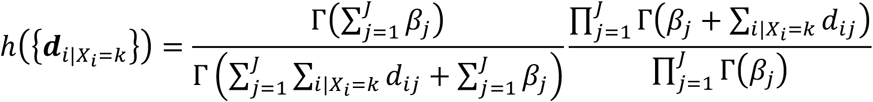

Formally, this model equates a resource niche (state) to a multinomial density and thereby assumes that resource use observations for taxa belonging to the same state are independent and identically distributed. As a model for count data it is closely related to topic models of word composition in a collection of text documents (Blei et al. 2003; Yin and Wang 2014) and to population genetic models of allele frequency composition in a set of populations (Pritchard et al. 2000). The key difference here is the specification of a prior model for ***X***. Because taxa are related, ***X*** is the outcome of evolution and individual *X_i_* are not independent.

We model evolution as a CTMC process where the rate of change is the same between all states (i.e. there is no evolutionary trend in the model) but varies independently among lineages. Our prior model for ***X*** is then the familiar phylogenetic likelihood function (Felsenstein 1981). Following Huelsenbeck et al. (2008), we treat lineage-specific rates of evolution as nuisance parameters drawn independently from a Gamma prior with shape parameter *α* and rate parameter fixed to 1. This model induces the same distribution on ***X*** as a model where the number of expected state changes along a branch is the same for all branches. Steel (2011) refers to this as the ultra-common mechanism model (UCM) to mark its contrast with the no-common mechanism model (Tuffley and Steel 1997) from which it derives. Concretely, this means that the transition probabilities used in the pruning algorithm to compute *p*(***X***) are equal to

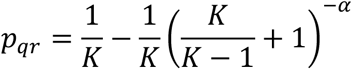

when ancestor and descendant states differ and to

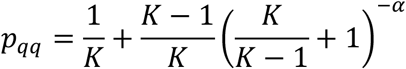

when ancestor and descendant states are the same. The full specification of the model is therefore

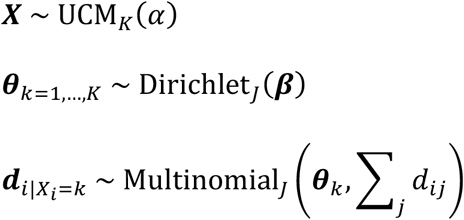

### Interpretation of hyperparameters

The hyperparameter *α* controls phylogenetic signal and equals the expected number of state changes along any branch separating ancestor and descendant. As *α* → 0, phylogenetic signal approaches 1 because descendants almost surely resemble their ancestors. As *α* → ∞, phylogenetic signal approaches 0 because a descendant’s state becomes independent of its ancestor’s state and resembles a random draw from a discrete uniform distribution. The hyperparameter ***β*** acts as a vector of pseudo-counts for each resource category. Larger values place more prior weight on observing a particular resource category in a set of counts than smaller values. This has implications for how the model makes decisions about when to separate taxa into different resource states. Differences among resource categories assigned a low prior weight contribute more toward separating taxa into distinct resource states than differences among resource categories assigned a high prior weight.

### Posterior inference

Inference under the model results in a sequence of samples of resource state assignments *X* for terminal taxa that approximate the posterior distribution *p*(***X***|***D***). Each sample partitions terminal taxa into some number *c*(***X***) ≤ *K* of groups, and each such group implies a posterior distribution over the parameter ***θ***_*k*_ of the multinomial density that the group represents. Specifically, due to the conjugacy of the multinomial and Dirichlet distributions the posterior distribution of each ***θ***_*k*_ is also Dirichlet with parameter {*β*_1_ + Σ_*i*|*X_i_*=*k*_ *d*_*i*1_,…,*β_j_* + Σ_*i*|*X_i_*=*k*_ *d*_*ij*_}. Therefore, inference of ***X*** implicitly yields posterior information about both the number of distinct resource niches and the relative importance of each resource category to each resource niche. Furthermore, because *p*(***X***) accounts for phylogenetic relatedness among terminal taxa, each sample of ***X*** induces a posterior distribution over ancestral resource states which we can explore using, e.g., stochastic character mapping (Nielsen 2002) or marginal ancestral state reconstructions (Schluter et al. 1997).

### Influence of K

We define phylogenetic signal as the quantity

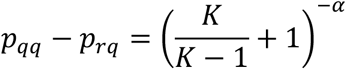

which ranges from 0 to 1 and quantifies how much information a descendant’s state provides about the state of its ancestor (Royer-Carenzi et al. 2013). A value of 1 means the ancestor was almost surely in the same state while a value of 0 means the ancestor was just as likely to have been in a different state than in the same state. Viewed as a function of *K*, phylogenetic signal approaches the curve 2^−*α*^ as *K* → ∞. Compared to a smaller value of *K*, a larger value of *K* therefore results in greater phylogenetic signal under the prior model for any given value of *α*. Because a large phylogenetic signal imposes a relatively greater prior penalty on events of character state change than a small phylogenetic signal, increasing *K* is expected to decrease the range of values of *c*(***X***) observed in posterior samples of ***X*** because closely related taxa will preferentially be placed into a shared resource state unless the data provide strong evidence against such a grouping. We note that the symmetry in the prior model means that the algorithmic complexity of evaluating *p*(***X***) is related to *c*(***X***) and not to *K*. This means that larger values of *K* may actually speed up computation relative to smaller values.

### Implementation

A Bayesian implementation of the model is available as an R package from github.com/blueraleigh/macroevolution. A Gibbs sampler for *p*(***X***|***D***) is derived using results from Yin and Wang (2014) and Schadt et al. (1998) and presented in the Appendix. The hyperparameters *α* and ***β*** are updated using slice sampling (Neal 2003).

### Simulation study

To illustrate application of the method we designed a simulation study using an empirical dataset on pseudoboine snake diets (Alencar et al. 2013). Our rationale for basing simulations on an empirical dataset is to ensure that properties of the data used to evaluate performance of the method are consistent with real studies, especially the distribution of observations per taxon and the distribution of resource specialization. Pseudoboine snakes are common members of the squamate reptile communities found in lowland rainforests of South America. Predominantly terrestrial or semi-arboreal, these snakes mainly eat small mammals, lizards, and other snakes. The dataset includes 606 observations of prey items from 8 prey categories for 32 species of pseudoboine snakes. Per species sample sizes range from 1 to 56 observations. We reanalyzed these data using a 33-species pseudoboine phylogeny extracted from the posterior distribution of trees in Tonini et al. (2016). We chose to use a different phylogeny from the original authors because their phylogeny excluded many species represented in the empirical dataset by small sample sizes, including some with the potential to impact state reconstructions at deeper nodes (Fig. 2). Using a more fully sampled phylogeny allowed us to better explore the influence of sample size on parameter estimation. The original publication coded each species with at least 8 diet observations into a set of 5 specialist diet states and 1 generalist diet state. Species were considered specialists if the prey resource represented at least 70 percent of recorded prey items (as in our Figure 1). When we applied the resampling procedure illustrated in Figure 1 to this empirical dataset under the assumption that the original state assignments represented the “truth”, we found that in approximately 20 percent of resampled datasets at least one original state (not always the same state) was not present and that in about 84 percent of cases at least one species was coded incorrectly (although overall coding accuracy was high, ranging from 0.77 to 1). Thus, this dataset illustrates some of the concerns raised in our introduction but is also well-sampled enough and shows enough variation to facilitate the estimation of separate multinomial distributions.

**Figure 2.**
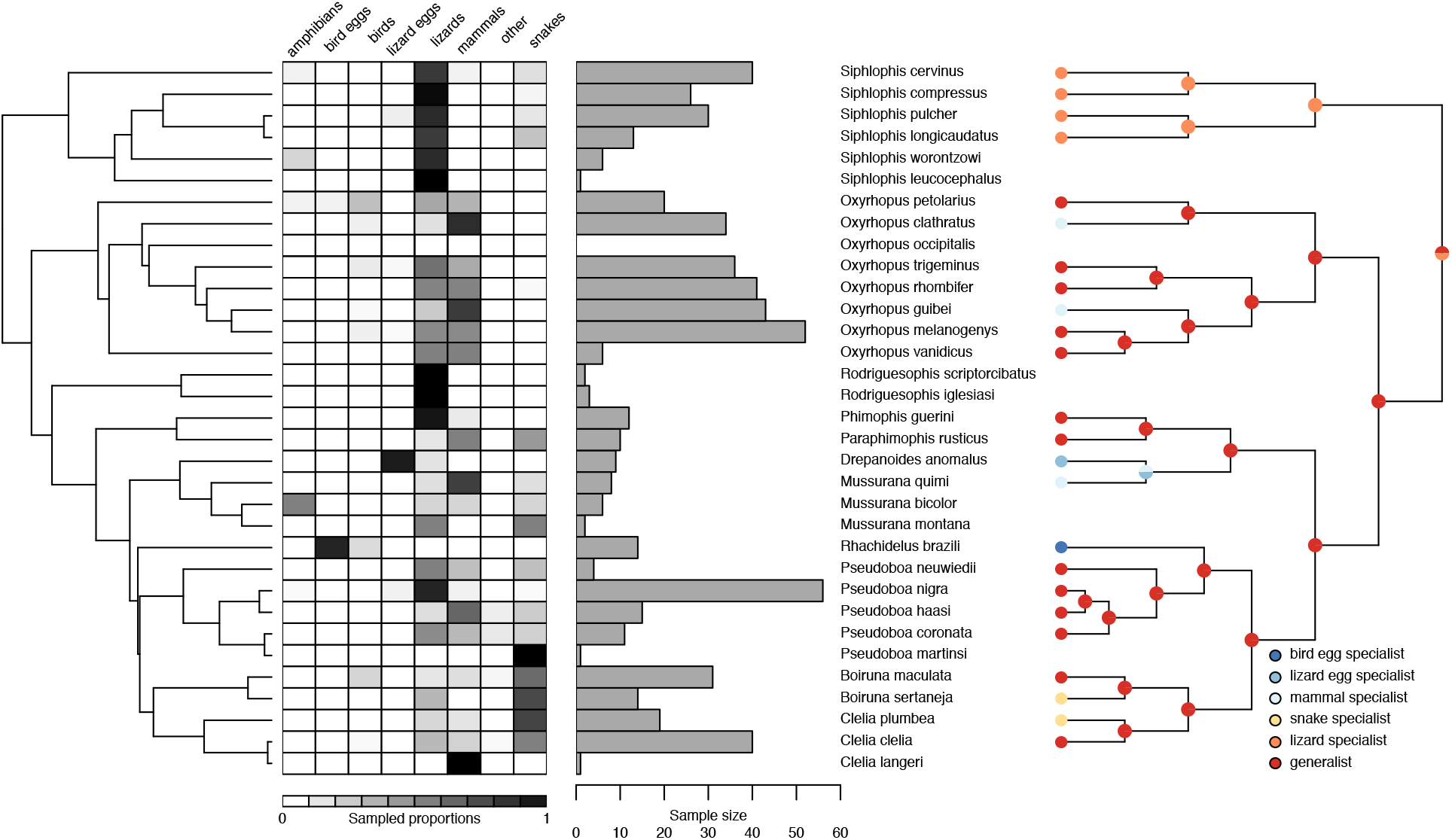
Summary attributes of the snake dietary dataset used to parameterize the simulation study, including phylogeny of pseudoboine snakes from Tonini et al. (2016) (left), and relative prey frequencies (middle left), total numbers of food observations per snake species (middle right), and cladogram with univariate ecological state encodings plus maximum parsimony reconstruction from Alencar et al. (2013) (right). Dark colors in the prey frequency matrix indicate higher sampled proportions of a particular prey item in a given diet. Our decision to use a different phylogeny from Alencar et al. (2013) was motivated by the observation that the original study excluded many species with low sample sizes, including species in phylogenetically informative positions with the potential to impact reconstructed evolutionary scenarios at deeper nodes (i.e., *Rodriguesophis*). In the present approach, ecological states are assumed to be multivariate probability distributions from which observed data are sampled, and the model estimates these distributions from the empirical data. In this case, even species with small sample sizes are informative and are therefore not excluded.

We first analyzed the empirical dataset assuming a prior model with *K* = 20 states, and we ran a Markov chain for 160000 iterations sampling every 128 iterations while holding the ***β*** hyperparameter at a constant value of {1,…, 1}. We subsequently extracted the maximum a posteriori estimate of ***X***, which implied a total of 6 resource states, and set the multinomial parameters for each of these states to their posterior mean value. We then simulated 20 datasets at each of 7 different levels of phylogenetic signal (0.1, 0.3, 0.5, 0.6, 0.7, 0.8, and 0.9) under the UCM model assuming *K* = 2, 3, 4, 5 and 6 resource states, and used the empirically estimated multinomial parameters and the empirical sample size distribution to generate prey resource use observations for each dataset. We analyzed the 700 simulated datasets assuming a prior model with *K* = 20 resource states, and we ran each Markov chain for 160000 iterations sampling every 128 iterations, again keeping the ***β*** hyperparameter at a constant value of {1,…, 1}.

To assess model performance, we compute a per-taxon accuracy index that measures how closely the multinomial density estimated for a taxon matches the multinomial density that generated the taxon’s data. Specifically, for terminal taxa accuracy is computed as

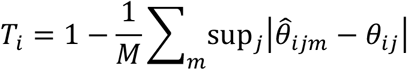

where 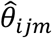 is the posterior mean estimate of the *j*-th multinomial proportion for the *i*-th taxon in the *m*-th posterior sample and *θ_ij_* is the true proportion and the average is taken over *M* posterior samples. The summand measures the largest absolute difference in proportions assigned by true and estimated multinomial densities to the same resource category. When the estimated and true multinomial densities perfectly coincide, the summand will equal 0 and accuracy will be 1. We can compute a similar metric for internal nodes using marginal ancestral state reconstructions as

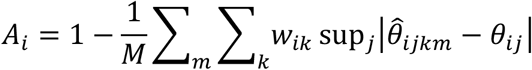

where *w_ik_* is the marginal posterior probability that the *i*-th internal node is assigned to resource state *k*.

## RESULTS

Results from the empirical dataset suggest at least five distinct trophic modalities among pseudoboine snakes with a posterior mode of six, corresponding to three highly specialized and three rather generalized diets (Fig. 3). Marginal posterior probabilities from ancestral state reconstructions reveal a scenario where a specialized diet of lizards was likely ancestral for the clade. Two subsequent dietary niche shifts occurred: one to a diet of lizards + mammals, and another to lizards + mammals + snakes. Interestingly, there is no evidence for a stepwise expansion of diet to first include mammals and then snakes. Rather, it appears these two states evolved independently: the ancestor of the lizard + mammal + snake diet is inferred to feed on lizards alone, rather than lizards + mammals. The lizard + mammal expansion occurred in the ancestor of the genus *Oxyrhopus*, and the lizard + mammal + snake expansion occurred in the ancestor of the clade containing *Pseudoboa* and *Mussurana*. Several lineages in this latter clade appear to have then undergone niche contractions to specialist diets of lizards (*Pseudoboa nigra*), lizard eggs (*Drepanoides anomalus*), and bird eggs (*Rhachidelus brazili*).

**Figure 3.**
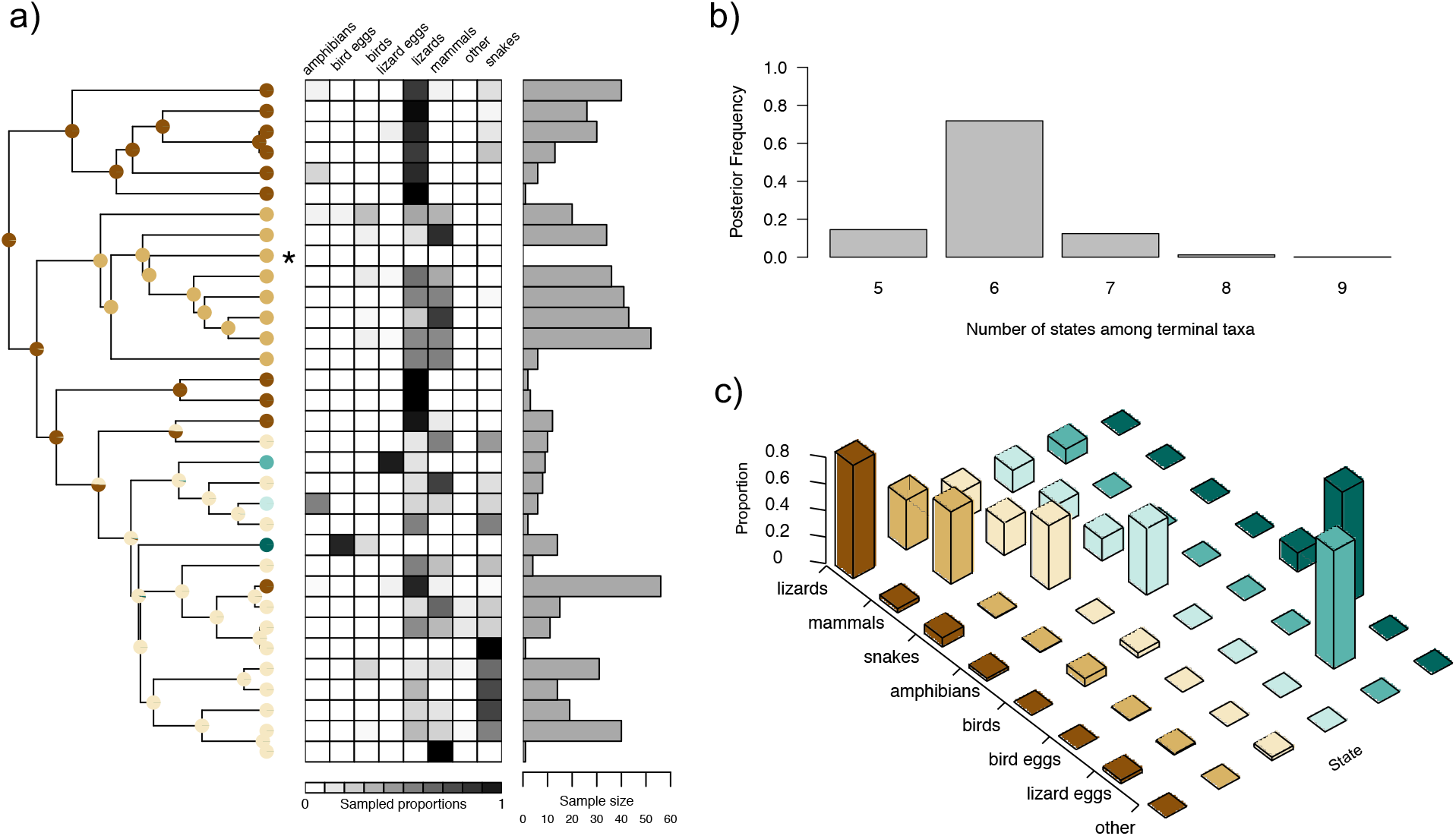
Reanalysis of the dataset presented in Figure 2 using the new model. a) The phylogeny and sampled diet observations were used to infer the diet states (colored circles) and their evolutionary history (marginal ancestral state probabilities shown as pie charts on the phylogeny). b) Posterior frequency distribution of the number of diet states among terminal taxa, computed from the posterior distribution of tip state assignments. The tip states in panel (a) depict the maximum a posteriori estimate from the posterior distribution of tip state assignments. Note that diet states are not observed directly, even at the tips of the tree; rather, all observed data are assumed to be sampled from a set of multinomial distributions. c) The maximum a posteriori estimate of the multinomial distributions for the diet states depicted on the phylogeny in panel (a). Here, the model inferred 6 states, corresponding to 3 specialist (> 70% specificity for a single prey group) and 3 generalist diets. Note that the terminal node marked with an asterisk is missing data, and information about its probable diet state is drawn only from what the model has learned about the states of its neighbors and the likelihood of evolutionary change.

Analysis of simulated datasets show that posterior frequency distributions of the number of resource states among terminal taxa were highly concentrated or centered on the number of resource states used to generate simulated datasets, indicating that the data were informative about the number of distinct resource use patterns among terminal taxa (Fig. 4). For taxa with moderate sample sizes, the posterior mean estimate of each taxon’s multinomial density was highly similar to the true distribution in the generating model (Fig. 5). This high similarity persisted across varying levels of phylogenetic signal, but small sample sizes resulted in lower accuracy when phylogenetic signal was low, creating in a distinctly triangular distribution of accuracy scores for terminal taxa. By contrast, accuracy scores for internal nodes decreased monotonically with decreasing phylogenetic signal and showed a close correspondence to the lower boundary observed for terminal node accuracy scores (Fig. 6). Because terminal nodes are associated with observational data, their states can be reasonably well-estimated even when phylogenetic signal is low. When signal is low, however, it is very difficult to estimate states at internal nodes because the observational data at tip nodes contain little information about ancestral states in this case.

**Figure 4.**
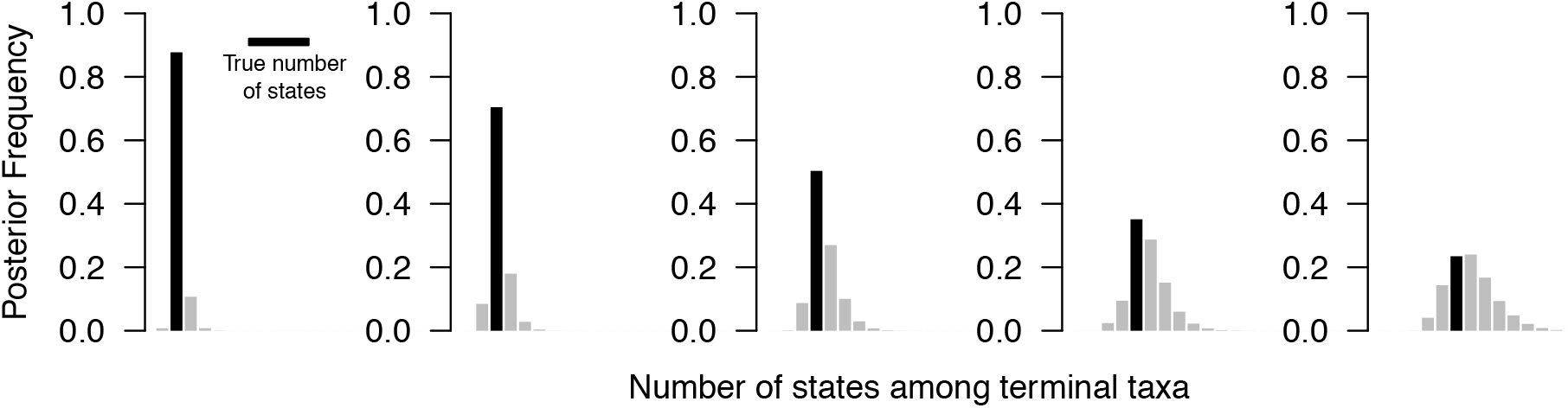
Each bar chart depicts the posterior frequency distribution of the number of resource states observed among the set of extant taxa; black bars highlight the number of resource states in the generating model. In all cases the prior model assumed there were 20 resource states that terminal taxa could potentially occupy.

**Figure 5.**
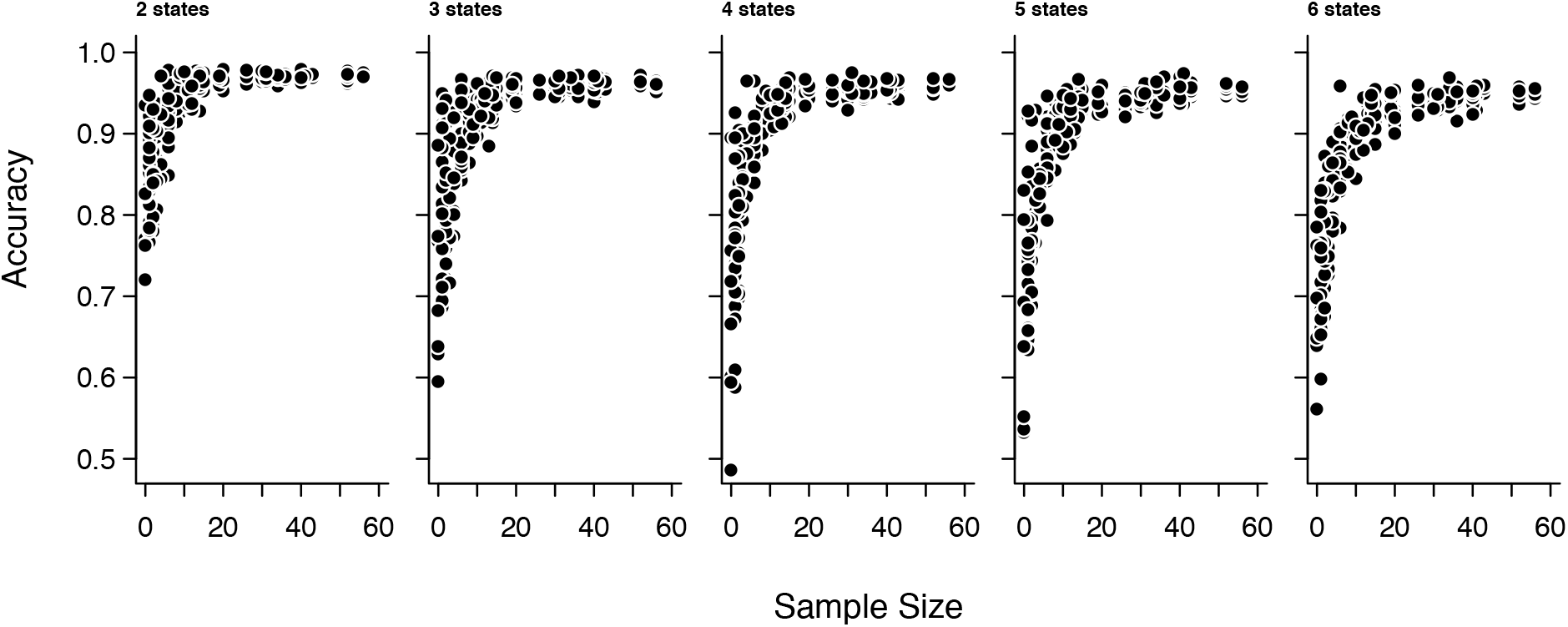
Accuracy of model-inferred resource use distributions for terminal taxa as a function of the number of resource use observations available. Each point corresponds to a terminal taxon in one simulation. Accuracy is a measure of the difference between true and estimated multinomial densities. An accuracy of 1 means that the posterior mean estimate of the multinomial density and the true multinomial density assign precisely the same proportions to each resource category. See main text for more details. Accuracy is consistently high across all simulations for taxa with moderate sample sizes.

**Figure 6.**
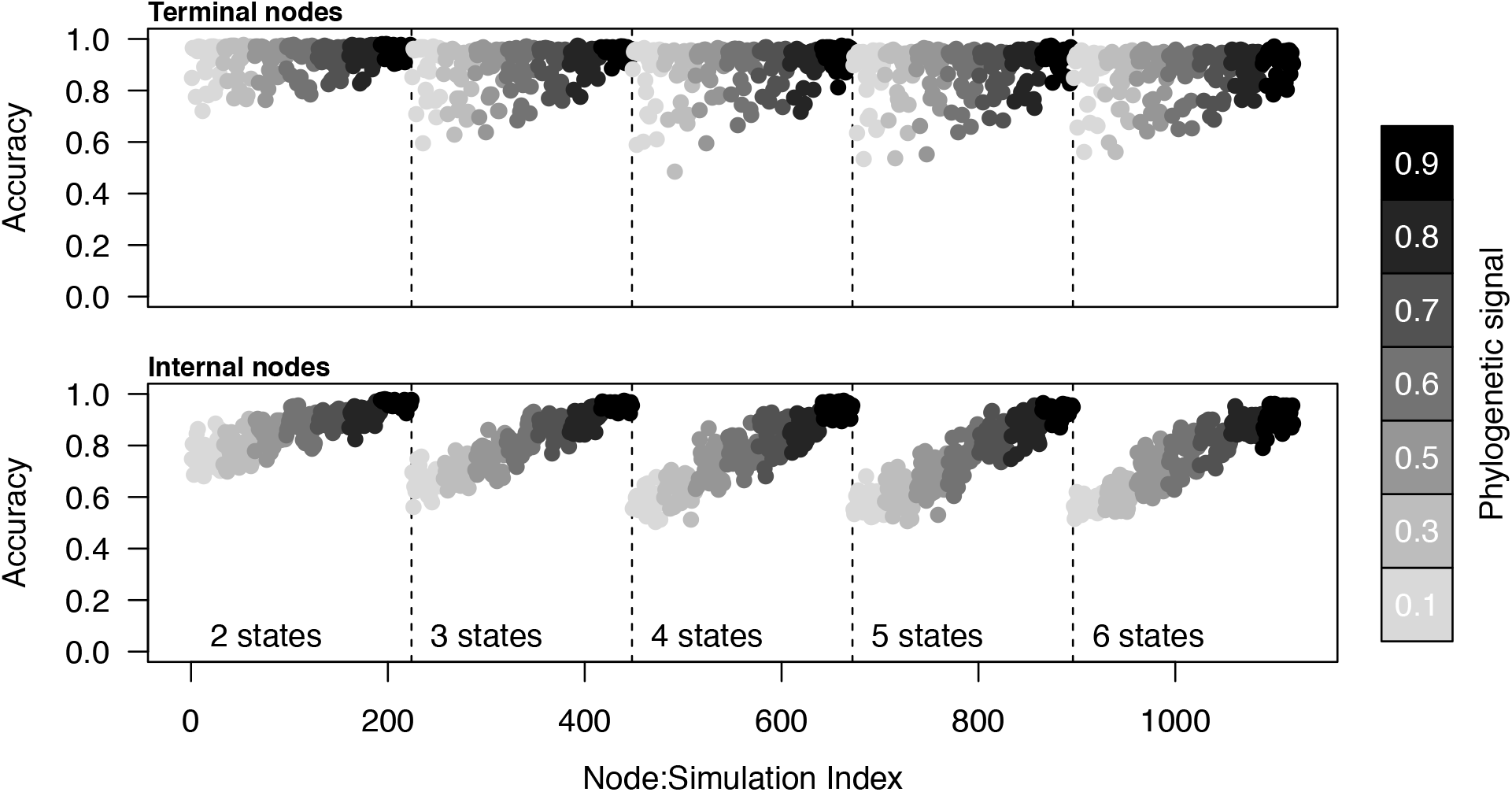
Accuracy of model-inferred resource use distributions for terminal taxa and internal nodes. Each point corresponds to a node in one simulation and points are colored according to the phylogenetic signal that was used to generate the simulated dataset. Accuracy is a measure of the difference between true and estimated multinomial densities. An accuracy of 1 means that the posterior mean estimate of the multinomial density and the true multinomial density assign precisely the same proportions to each resource category. In the case of internal nodes, differences are also averaged using the marginal posterior probabilities assigned to the different resource use distributions. See main text for more details. The triangular distribution of accuracy for terminal nodes is evidently caused by small sample sizes (cf. Fig. 4). Accuracy for internal nodes decreases monotonically with decreasing phylogenetic signal and appears to track the lower bound for accuracy of terminal taxa. Because tip nodes are associated with observational data, their states can be estimated with reasonable accuracy even when phylogenetic signal is low. However, low signal makes it very difficult to reconstruct internal node states, as expected because tip states provide very little information about the states of their ancestors in this case.

## DISCUSSION

We developed a comparative method for macroevolutionary analysis of multivariate count data. The method is general and may be applied to any data expressible as a set of observational counts from different categories. Such datatypes arise frequently in community ecology and behavior. Potential applications include the comparative analysis of diet, foraging behavior, activity patterns, and habitat preferences. The method is similar to standard continuous-time Markov chain (CTMC) models of phenotypic evolution but differs in several important respects. First, the number of states in the model and the states themselves are unobserved and must be estimated from empirical data on resource use. Second, each state is a multinomial distribution rather than a categorical variable. This latter property enables researchers to model complex multidimensional phenotypes that cannot readily be expressed in univariate space without considerable loss of information. Moreover, this property allows the method to accommodate uncertainty in the state assignments of terminal taxa that arises from the effects of sampling variation.

Simulations revealed that the new method is generally able to determine the correct number of states and that it provides accurate estimates of the underlying multinomial distributions, both for terminal taxa as well as internal nodes. We designed simulations around empirical patterns of resource use in a dataset on snake diets (Alencar et al. 2013). Therefore, caution is warranted in generalizing the good performance observed in the current study to other datasets. In particular, performance of the model will depend on the idiosyncrasies of individual datasets, including the distribution of sample sizes and the distribution of overlaps in resource use among species. We expect that states represented by few observations will be difficult to infer, especially if those states show appreciable overlap with other states.

Our empirical analysis identified at least two feeding modalities among the set of species Alencar et al. (2013) recognize as “generalists”: species that feed predominantly on snakes but that regularly eat lizards and mammals, and species that feed predominantly on mammals and lizards. Ancestral state estimates strongly suggest that each of these feeding modalities arose from a more specialized diet comprised almost entirely of lizards. This is in contrast to the results of Alencar et al. (2013), which imply that nearly all origins of specialized feeding modalities occurred from a generalist ancestor. Interestingly, the model finds evidence that the data-poor species *Mussurana bicolor* occupies a distinct dietary niche (cf. Fig. 3). This is because the relatively high proportion of amphibians in *M. bicolor* diet samples, although supported by few observations, is highly unlikely given the complete absence of amphibian prey items from its close relatives. However, because *M. bicolor* has such a poorly characterized diet a more conservative prior model (larger *K*) can easily cause *M. bicolor* to fall back into the state of its close relatives. The model also fails to recognize the small mammal and snake “specialist” states recognized by Alencar et al. (2013) even though the species representing these states have larger sample sizes than *M. bicolor*. Our explanation for this is that both mammals and snakes are commonly observed prey items in the diets of species closely related to these putative mammal and snake specialists. As a result, the amount of data needed to recognize a specialist on one of these prey resources is substantially greater than what would be needed to recognize a specialist on a prey resource that is rarely or never observed in the diet of close relatives.

As currently implemented, the approach described here does not directly model evolutionary gains and losses or substitutions of different resources. Indeed, under the prior model no resource is ever truly absent from the reconstructed states (although its proportional representation may approach zero as values in ***β*** become small). This contrasts with biogeographic-type models that explicitly model resource use expansions, contractions, and substitutions (e.g. see Hardy (2017) for application of Ree and Smith’s (2008) dispersal-extinction-cladogenesis model to binary encoded diet data). Although these types of changes are implicit in the sequence of reconstructed states derived from the model, future studies might want to explore how to combine more complex evolutionary models with the current model for count data. Nonetheless, the advantage of a simple evolutionary model is that it has broad scope. It would be possible, for example, to apply the current approach to continuous characters by keeping the same evolutionary model but changing the model for observations from a multinomial to a multivariate-normal distribution, which could then be applied to other data types used for quantifying resource use such as stable isotope ratios of carbon and nitrogen.

One challenge for comparative methods is their limited ability to model ecological phenotypes that cannot be neatly summarized by a single value (Hardy & Linder 2005; Hardy 2006). Recent years have seen progress in this direction for continuous traits, including models that accommodate intraspecific variation, function-valued traits, and other non-gaussian data (Ives et al. 2007; Felsenstein 2008; Evans et al. 2009; Jones & Moriarty 2013; Goolsby 2015; Quintero et al. 2015). The general approach developed here, where each state is a multinomial distribution rather than a categorical variable, extends this progress to traits like diet and habitat that are typically treated as univariate categorical variables. By placing an emphasis on individual natural history observations, the method draws attention to the central role such observations play in evolutionary biology (Greene 1986) and to the many remaining opportunities for developing comprehensive ecological databases that advance our understanding of biodiversity (Hortal et al. 2015).

## SUMMARY

We described a novel methodological framework for studying the evolutionary dynamics of complex ecological traits on phylogenetic trees. Previous approaches to this problem have assumed that ecological states are categorical variables and that species are monomorphic for particular states. We relaxed this assumption through the use of a CTMC model that treats ecological states as probability distributions from which observed data are sampled. Results from our model provide a much richer understanding of macroevolutionary patterns than past approaches and can help illuminate the “phylogenetic natural history” of particular systems (Uyeda et al. 2018). Although our method is designed for the analysis of multivariate count data, we suggest that the approach of treating states as probability distributions has wide applicability and will greatly facilitate the comparative analysis of novel sources of ecological data.

## ACKNOWLEDGEMENTS

This research was supported in part through computational resources and services provided by Advanced Research Computing at the University of Michigan, Ann Arbor. This material is based upon work supported by the National Science Foundation Graduate Research Fellowship and by the University of Michigan Department of Ecology and Evolutionary Biology Block Grant. We thank the Associate Editor and two anonymous reviewers whose comments helped improve the manuscript.

## APPENDIX

If we update ***X*** one terminal taxon at a time by sampling a new state from the proposal distribution *p*(*X*_*i*_ = *k*|***X***_−*i*_,**D**), the Metropolis-Hastings acceptance ratio will equal 1. We derive this distribution below using results from Yin and Wang (2014) and Schadt et al. (1998) and present an algorithm for its implementation. First, we have

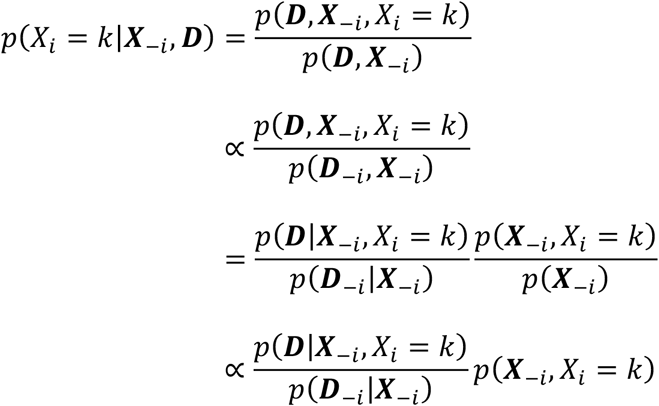

The symbols ***D***_−*i*_ and ***X***_−*i*_ is refer to the data matrix and resource state vector with terminal taxon *i* excluded. The first proportionality follows because *p*(***D***,***X***_−*i*_) can be written as *p*(***d***_*i*_,***D***_−*i*_,***X***_−*i*_) = *p*(***D***_−*i*_,***X***_−*i*_|***d***_*i*_)p(***d_i_***). But ***d**_i_* contains no information about (***D***_−*i*_, ***X**_−*i*_)* so *p*(***D**_−i_*,***X***_−*i*_|***d**_i_*)=*p*(***D***_−*i*_, ***X***_−*i*_) and we simply ignore the constant *p*(***d**_i_*). The second proportionality follows because *p*(***X**_-i_*) does not depend on *i* and hence is the same for all values of *X_i_*. Yin and Wang (2014) derive 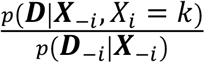 which can be written as

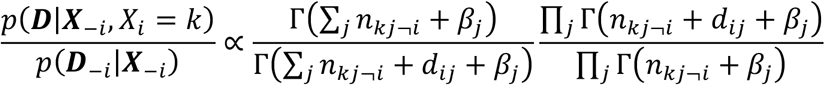

where *n_kj_* = Σ_*h*|*X_h_* =*k*_ *d_hj_* and *n*_*kj*¬*i*_ = *n*_*kj*_ – *d_ij_*. To compute *p*(***X***_−*i*_, *X_i_* = *k*) we make use of the recurrence relations described Schadt et al. (1998). Let *a*(*m*) denote *m*’s ancestor and let *s*(*m*) denote m’s sibling. We write *s*(*m*)=*r* and *s*(*m*)=*l* when *m* is a left, respectively right, descendant *a*(*m*). With ***T*** representing the entire phylogeny, we let ***T**_m_* denote the subtree rooted at node *m* and we let ***T**_−m_* denote the complementary subtree that results from pruning ***T**_m_* from ***T***. The notation {*X_h_*, *h* ∈ ***T**_m_* is used to indicate the set of resource states in ***X*** where *h* is a terminal node in ***T**_m_*. We implicitly augment the state vector ***X*** to 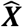 so that it include states at terminal and internal nodes and use the notation 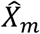 to refer to the state of any node. If *m* is a terminal node 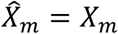. Let ***F**_m_* = {*F_m_*(1),…, *F_m_*(*K*)} where each 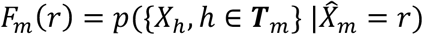 and let ***S**_m_* = {*S_m_*(1),…,*S_m_*(*K*)} where each *S_m_*(*q*) = Σ_r_ *P*_*qr*_*F_m_*(*r*) and *p_qr_* denotes the transition probability. ***F**_m_* and ***S**_m_* are computed with a postorder traversal of ***T*** using the standard peeling algorithm (Felsenstein 1981). Now let ***U**_m_* = {*U_m_*(1),…, *U_m_*(*K*)} where each 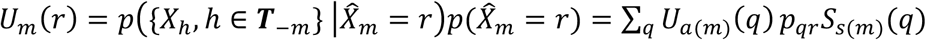. ***U**_m_* can be computed with a preorder traversal of ***T*** after defining 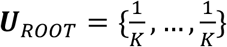. When *m* is a terminal node the joint probability 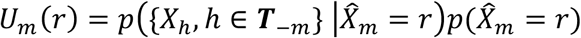 can be written as *U_m_*(*r*) = *p*(***X***_−*m*_ | *X_m_* = *r*)*p*(*X_m_* = *r*) = *p*(*X_m_* = *r*, ***X***_−*m*_). Putting these threads together gives

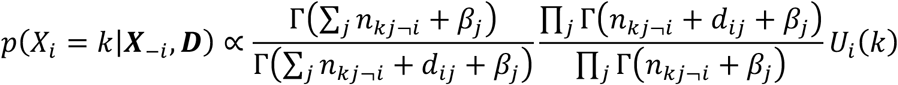

With ***F***, ***S***, and ***U*** precomputed from an initial configuration of ***X*** we use algorithm A.1 to update ***X***, which assumes each internal node has only two immediate descendants and that a preorder traversal always visits a left descendant before visiting a right descendant.

### Algorithm A. 1

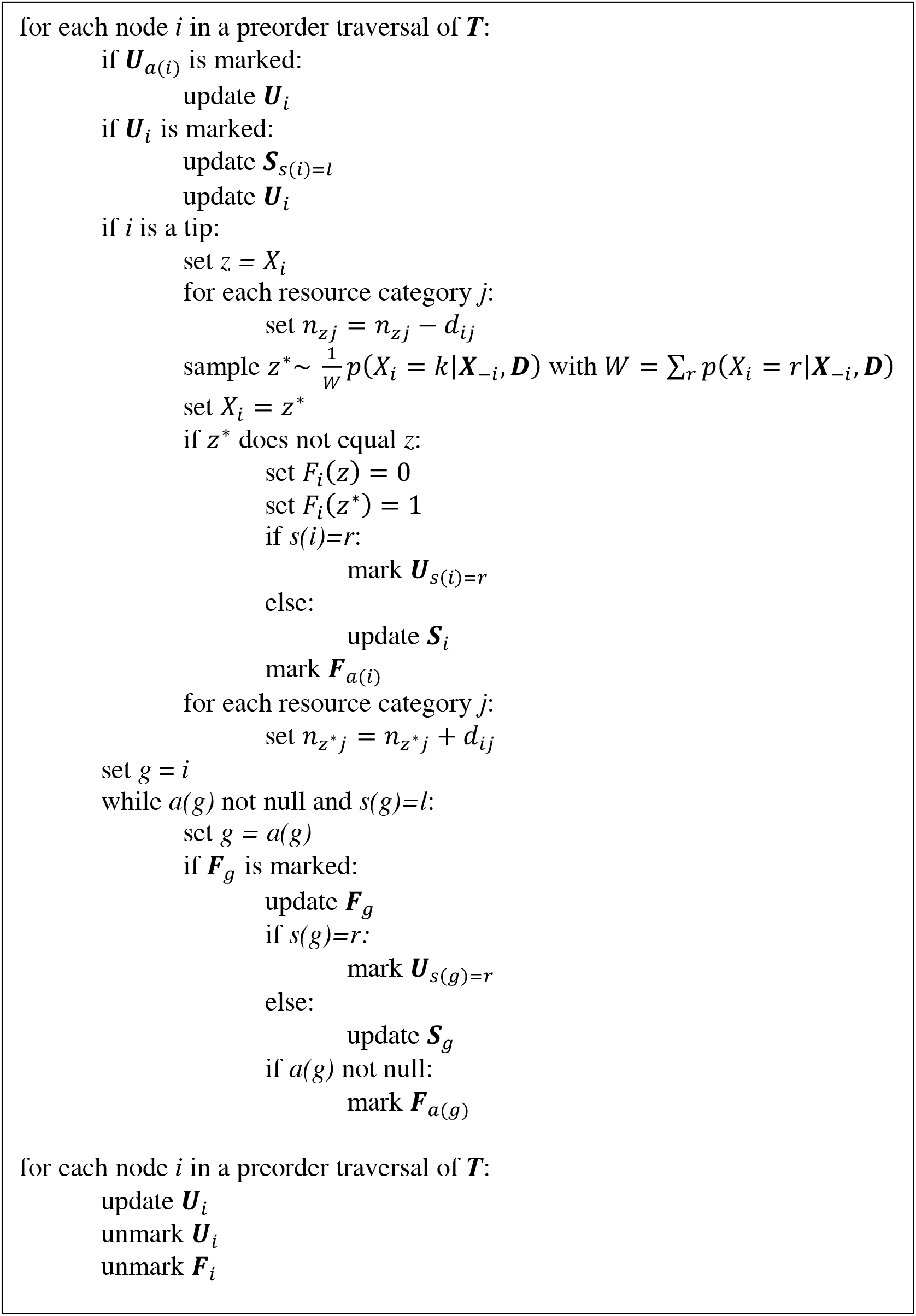

Algorithm A.1 ensures that whenever we visit a terminal node the local ***U**_i_* needed to compute *p*(*X_i_* = *k*|***X***_−*i*_, ***D***) are correct. When the first preorder traversal of ***T*** has finished ***F*** and ***S*** correctly specify the conditional likelihood arrays for the new ***X***. However, the update operations leave the joint likelihood arrays in ***U*** out of date even though they are locally correct for each individual update. Therefore, a second preorder traversal of ***T*** to update ***U*** is necessary before updating ***X*** again.

## LITERATURE CITED

Alencar L.R.V., Gaiarsa M.P., Martins M. 2013. The evolution of diet and microhabitat use in pseudoboine snakes. South American Journal of Herpetology. 8:60–66.

Bassar R.D., Marshall M.C., López-Sepulcre A., Zandona E., Auer S.K., Travis J., Pringle C.M., Flecker A.S., Thomas S.A., Fraser D.F., Reznick D.N. 2010. Local adaptation in Trinidadian guppies alters ecosystem processes. PNAS. 107:3616–3621.

Baum L.E., Petrie T. 1966. Statistical Inference for Probabilistic Functions of Finite State Markov Chains. The Annals of Mathematical Statistics. 37:1554–1563.

Beaulieu J.M., O’Meara B.C. 2016. Detecting hidden diversification shifts in models of trait-dependent speciation and extinction. Systematic Biology. 65:583–601.

Beaulieu J.M., O’Meara B.C., Donoghue M.J. 2013. Identifying hidden rate changes in the evolution of a binary morphological character: the evolution of plant habit in Campanulid angiosperms. Systematic Biology. 62:725–737.

Blei D.M., Ng A.Y., Jordan M.I. 2003. Latent Dirichlet Allocation. Journal of Machine Learning Research. 3:993–1022.

Burin G., Kissling W.D., Guimarães P.R., Sekercioglu C.H., Quental T.B. Omnivory in birds is a macroevolutionary sink. Nature Communications. 7:11250.

Caetano D.S., Beaulieu J.M., O’Meara B.C. 2018. Hidden state models improve state-dependent diversification approaches, including biogeographical models. Evolution. 72:2308–2324.

Cantalapiedra J.L., FitzJohn R.G., Kuhn T.S., Hernández Fernández M., DeMiguel D., Azanza B., Morales J., Mooers A.Ø. 2014. Dietary innovations spurred the diversification of ruminants during the Caenozoic. Proceedings of the Royal Society B: Biological Sciences. 281:20132746.

Davis A.M., Unmack P.J., Vari R.P., Betancur-R R. 2016. Herbivory promotes dental disparification and macroevolutionary dynamics in grunters (Teleostei: Terapontidae), a freshwater adaptive radiation. The American Naturalist. 187:320–333.

Ehrlich P.R., Raven P.H. 1964. Butterflies and plants: a study in coevolution. Evolution. 18:586–608.

Evans M.E.K., Smith S.A., Flynn R.S., Donoghue M.J. 2009. Climate, niche evolution, and diversification of the “bird-cage” evening primroses (*Oenothera*, Sections *Anogra* and *Kleinia*). The American Naturalist. 173:225–240.

Felsenstein J. 1981. Evolutionary trees from DNA sequences: A maximum likelihood approach. J Mol Evol. 17:368–376.

Felsenstein J. 2008. Comparative methods with sampling error and within-species variation: contrasts revisited and revised. The American Naturalist. 171:713–725.

Felsenstein J. 2012. A comparative method for both discrete and continuous characters using the threshold model. The American Naturalist. 179:145–156.

Felsenstein, J., Churchill G.A. 1996. A Hidden Markov Model approach to variation among sites in rate of evolution. Molecular biology and evolution. 13:93–104.

Forister M.L., Dyer L.A., Singer M.S., Stireman III J.O., Lill J.T. 2012. Revisiting the evolution of ecological specialization, with emphasis on insect-plant interactions. Ecology. 93:981–991.

Futuyma D.J., Moreno G. 1988. The evolution of ecological specialization. Annual Review of Ecology and Systematics. 19:207–233.

Gillespie R. 2004. Community assembly through adaptive radiation in Hawaiian spiders. Science. 303:356–359.

Givnish T.J., Barfuss M.H.J., Van Ee B., Riina R., Schulte K., Horres R., Gonsiska P.A., Jabaily R.S., Crayn D.M., Smith J.A., Winter K., Brown G.K., Evans T.M., Holst B.K., Luther H., Till W., Zizka G., Berry P.E., Sytsma K.J. 2014. Adaptive radiation, correlated and contingent evolution, and net diversification in Bromeliaceae. Molecular Phylogenetics and Evolution. 71:55–78.

Goolsby E.W. 2015. Phylogenetic Comparative Methods for Evaluating the Evolutionary History of Function-Valued Traits. Systematic Biology. 64:568–578.

Greene H.W. 1986. Natural history and evolutionary biology. In: Feder M.E., Lauder G.V., editors. Predator–prey relationships: perspectives and approaches from the study of lower vertebrates. University of Chicago Press. p. 198.

Hardy C.R. 2006. Reconstructing ancestral ecologies: challenges and possible solutions. Diversity and Distributions. 12:7–19.

Hardy C.R., Linder H.P. 2005. Intraspecific variability and timing in ancestral ecology reconstruction: a test case from the cape flora. Systematic Biology. 54:299–316.

Hardy N.B. 2017. Do plant-eating insect lineages pass through phases of host-use generalism during speciation and host switching? Phylogenetic evidence. Evolution. 71:2100–2109.

Hardy N.B., Otto S.P. 2014. Specialization and generalization in the diversification of phytophagous insects: tests of the musical chairs and oscillation hypotheses. Proceedings of the Royal Society B: Biological Sciences. 281:20132960.

Harmon, L. J., Matthews B., Des Roches S., Chase J.M., Shurin J.B., Schluter D. 2009. Evolutionary diversification in stickleback affects ecosystem functioning. Nature. 458:1167–1170.

Hortal J., de Bello F., Diniz-Filho J.A.F., Lewinsohn T.M., Lobo J.M., Ladle R.J. 2015. Seven shortfalls that beset large-scale knowledge of biodiversity. Annual Review of Ecology, Evolution, and Systematics. 46:523–549.

Huelsenbeck J.P., Ané C., Larget B., Ronquist F. 2008. A Bayesian perspective on a non-parsimonious parsimony model. Systematic Biology. 57:406–419.

Ives A.R., Midford P.E., Garland Jr T. 2007. Within-species variation and measurement error in phylogenetic comparative methods. Systematic Biology. 56:252–270.

Jones N.S., Moriarty J. 2013. Evolutionary inference or function-valued traits: Gaussian process regression on phylogenies. Journal of the Royal Society Interface. 10:20120616.

Kelley S.T., Farrell B.D. 1998. Is specialization a dead end? The phylogeny of host use in *Dendroctonus* bark beetles (Scolytidae). Evolution. 52:1731–1743.

Losos J.B., Leal M., Glor R.E., de Queiroz K., Hertz P.E., Rodríguez Schettino L., Chamizo Lara A., Jackman T.R., Larson A. 2003. Niche lability in the evolution of a Caribbean lizard community. Nature. 424:542–545.

MacArthur R.H. 1958. Population ecology of some warblers of northeastern coniferous forests. Ecology. 39:599–619.

Marazzi B., Ané C., Simon M.F., Delgado-Salinas A., Luckow M., Sanderson M.J. 2012. Locating evolutionary precursors on a phylogenetic tree. Evolution. 66:3918–3930.

Martin C.H., Wainwright P.C. 2011. Trophic novelty is linked to exceptional rates of morphological diversification in two adaptive radiations of *Cyprinodon* pupfish. Evolution. 65:2197–2212.

Miller E.T., Wagner S.K., Harmon L.J., Ricklefs R.E. 2017. Radiating despite a lack of character: ecological divergence among closely related, morphologically similar honeyeaters (Aves: Meliphagidae) co-occurring in arid Australian environments. The American Naturalist. 189:E14–E30.

Mitter C., Farrell B., Wiegmann B. 1988. The phylogenetic study of adaptive zones: has phytophagy promoted insect diversification? The American Naturalist. 132:107–128.

Neal, Radford M. 2003. Slice Sampling. The Annals of Statistics. 31:705–767.

Nielsen R. 2002. Mapping mutations on phylogenies. Systematic Biology. 51:729–739.

Nosil P. 2002. Transition rates between specialization and generalization in phytophagous insects. Evolution. 56:1701–1706.

O’Meara B.C. 2012. Evolutionary inferences from phylogenies: a review of methods. Annual Review of Ecology and Systematics. 43:267–285.

Price S.A., Hopkins S.S.B., Smith K.K., Roth V.L. 2012. Tempo of trophic evolution and its impact on mammalian diversification. PNAS. 109:7008–7012.

Pritchard J.K., Stephens M., Donnelly P. 2000. Inference of Population Structure Using Multilocus Genotype Data. Genetics. 155:945–959.

Quintero I., Keil P.K., Jetz W., Crawford F.W. 2015. Historical biogeography using species geographical ranges. Systematic Biology. 64:1059–1073.

Ree R.H., Smith S.A. 2008. Maximum likelihood inference of geographic range evolution by dispersal, local extinction, and cladogenesis. Systematic Biology. 57:4–14.

Revell L.J. 2014. Ancestral character estimation under the threshold model from quantitative genetics. Evolution. 68:743–759.

Robinson G.S. 1999. HOSTS - a database of the hostplants of the world’s Lepidoptera. Nota Lepidopterologica. 22:35–47.

Royer-Carenzi M., Pontarotti P., Didier G. 2013. Choosing the best ancestral character state reconstruction method. Mathematical Biosciences. 242:95–109.

Schadt E.E., Sinsheimer J.S., Lange K. 1998. Computational Advances in Maximum Likelihood Methods for Molecular Phylogeny. Genome Res. 8:222–233.

Schluter D., Price T., Mooers A.Ø., Ludwig D. 1997. Likelihood of ancestor states in adaptive radiation. Evolution. 51:1699–1711.

Shine R. 1994. Allometric patterns in the ecology of Australian snakes. Copeia. 1994:851–867.

Steel M. 2011. Can we avoid “SIN” in the house of “no common mechanism”? Systematic Biology. 60:96–109.

Tonini J., Beard K.H., Barbosa Ferreira R., Jetz W., Pyron R.A. 2016. Fully-sampled phylogenies of squamates reveal evolutionary patterns in threat status. Biological Conservation. 204:23–31.

Tuffley C., Steel M. 1997. Links between maximum likelihood and maximum parsimony under a simple model of site substitution. Bulletin of Mathematical Biology. 59:581–607.

Uyeda J.C., Zenil-Ferguson R., Pennell M.W. 2018. Rethinking phylogenetic comparative methods. Systematic Biology. 67:1091–1109.

Vitt L.J., Vangilder L.D. 1983. Ecology of a snake community in northeastern Brazil. Amphibia-Reptilia. 4:273–296.

Vrba E.S. 1987. Ecology in relation to speciation rates: some case histories of Miocene-Recent mammal clades. Evolutionary Ecology. 1:283–300.

Yin J., Wang J. 2014. A dirichlet multinomial mixture model-based approach for short text clustering. Proceedings of the 20th ACM SIGKDD international conference on Knowledge discovery and data mining - KDD ‘ 14.:233–242.

